# *Hif-2α* programmes oxygen chemosensitivity in chromaffin cells

**DOI:** 10.1101/2023.07.14.548982

**Authors:** Maria Prange-Barczynska, Holly A. Jones, Yoichiro Sugimoto, Xiaotong Cheng, Joanna D. C. C. Lima, Indrika Ratnayaka, Gillian Douglas, Keith J. Buckler, Peter J. Ratcliffe, Thomas P. Keeley, Tammie Bishop

**Author notes:** **Correspondence to:** Peter J. Ratcliffe, Thomas P. Keeley and Tammie Bishop, Target Discovery Institute, Roosevelt Drive, Oxford, OX3 7FZ, UK. **Authorship note:** MPB, HAJ and YS are joint first authors. PJR, TPK and TB are co-senior authors.

## Abstract

The study of transcription factors that determine specialised neuronal functions has provided invaluable insights into the physiology of the nervous system. Peripheral chemoreceptors are neurone-like electro-physiologically excitable cells that link the oxygen content of arterial blood to the neuronal control of breathing. In the adult, this oxygen chemosensitivity is exemplified by the Type I cells of the carotid body and recent work has revealed one isoform of the transcription factor HIF, HIF-2α, to have a non-redundant role in the development and function of that organ. Here we show that the activation of HIF-2α, including isolated overexpression alone, is sufficient to induce oxygen chemosensitivity in the otherwise unresponsive adult adrenal medulla. This phenotypic change in the adrenal medulla was associated with retention of extra-adrenal paraganglioma-like tissues that resemble the foetal organ of Zuckerkandl and also manifest oxygen chemosensitivity. Acquisition of chemosensitivity was associated with changes in the adrenal medullary expression of classes of genes that are ordinarily characteristic of the carotid body, including G-protein regulators and atypical subunits of mitochondrial cytochrome oxidase. Overall, the findings suggest that, at least in certain tissues, HIF-2α acts as a phenotypic driver for cells that display oxygen chemosensitivity, providing a route to mechanistic understanding.

## Introduction

One of the most vital, but still incompletely understood, functions in homeostatic physiology is that of the arterial chemoreceptors that control breathing in response to changes in blood oxygen, carbon dioxide and other stresses (Weir et al. 2005; Semenza and Prabhakar 2018; Ortega-Saenz and Lopez-Barneo 2020; Holmes et al. 2022). This chemosensitivity is generally considered to be a restricted property of specialised cells within the central nervous system and the Type I cells of the carotid body, often termed peripheral chemoreceptors (Ortega-Saenz and Lopez-Barneo 2020). Carotid body Type I cells respond to low arterial oxygen by K^+^ channel inhibition, resulting in membrane depolarisation, voltage gated Ca^2+^ influx and dense core vesicle release to excite afferent neurones and stimulate respiration. Nevertheless, despite decades of intense study, the mechanisms that generate and transduce these oxygen-sensitive signals are incompletely understood (Weir et al. 2005; Semenza and Prabhakar 2018; Ortega-Saenz and Lopez-Barneo 2020; Holmes et al. 2022).

More recently, a new route to the understanding of these processes has been opened by the elucidation of oxygen sensing mechanisms that transduce transcriptional responses to changes in oxygen availability via hypoxia inducible transcription factors (HIFs)(Bishop and Ratcliffe 2014; Semenza and Prabhakar 2018). HIF is an α/β heterodimeric transcription factor, whose α subunit is hydroxylated in the presence of oxygen by HIF prolyl hydroxylases to target it for proteasomal degradation via the von Hippel-Lindau ubiquitin E3 ligase. In hypoxia, HIF-α subunits escape degradation and dimerise with HIF-β to activate an extensive transcriptional cascade that adapts cells to hypoxia. In keeping with the importance of oxygen homeostasis to animal life, all animals possess at least one HIF-PHD-VHL triad. However, gene duplication events at the base of vertebrate evolution have generated multiple HIF-α and HIF prolyl hydroxylase isoforms (Loenarz et al. 2011). Thus, mammalian species generally possess three HIF prolyl hydroxylases (termed prolyl hydroxylase domain, PHD 1, 2 and 3) and three HIF-α isoforms (HIF-1α, HIF-2α and HIF-3α).

Of particular interest is the HIF-2α isoform, which has been extensively characterised, and which shows features consistent with being a more ‘modern’ isoform connected to the function of specialised oxygen delivery organs such as lung ventilation and blood vascular systems that are found in higher animals (Tian et al. 1998; Wiesener et al. 2003; Cowburn et al. 2016). To date, one of the most striking of these connections is to the oxygen chemosensitive Type I cells of the carotid body. HIF-2α is expressed at uniquely high levels in Type I cells of the carotid body and genetic ablation of HIF-2α has revealed essential functions in carotid body development and in carotid body oxygen chemosensitivity, including the augmentation of hypoxic ventilatory response after acclimatisation to hypoxia (Tian et al. 1998; Hodson et al. 2016; Zhou et al. 2016; Gao et al. 2017; Fielding et al. 2018; Macias et al. 2018; Cheng et al. 2020; Moreno-Dominguez et al. 2020).

In the present work, we sought to determine whether the expression of HIF-2α might be sufficient to induce oxygen chemosensitivity in an organ that is otherwise insensitive. Here we show that overexpression of HIF-2α, either by inactivation of the principal HIF prolyl hydroxylase (PHD2) or by forced expression of HIF-2α alone, is sufficient to induce chemosensitivity in the adult adrenal medulla, which is not normally chemosensitive (Thompson et al. 1997). The induction of chemosensitivity in the adrenal medulla was associated with bidirectional changes in the expression of genes that are ordinarily more, or less, strongly expressed in the carotid body versus the adrenal medulla. Alterations in the adult adrenal medulla were accompanied by the retention of the organ of Zuckerkandl, which normally atrophies after birth (Coupland 1965), and ectopic adrenal medullary tissue, both of which also manifested oxygen chemosensitivity. Given reports of oxygen chemosensitivity in the foetal adrenal gland (Seidler and Slotkin 1985; Mojet et al. 1997; Thompson et al. 1997), these findings are consistent with the maintenance or induction of a foetal-like state in adrenal medulla and related tissues. The sufficiency of HIF-2α alone to drive these phenotypes suggests a direct impact on the physiology of oxygen chemosensitivity that should be mechanistically informative.

## Results

### Phd2 inactivation in the adrenal medulla induces carotid body enriched gene expression

With the aim of better understanding oxygen homeostasis, substantial effort has focussed on defining interactions between transcriptional signalling by the HIF pathway and oxygen chemosensitivity in the carotid body. As part of this work, we observed that inactivation of the principal oxygen sensor that regulates HIF (PHD2) resulted in paraganglioma-like changes in the carotid body, adrenal medulla and related tissues (Fielding et al. 2018; Eckardt et al. 2021). To pursue this further, we intercrossed mice to obtain *Phd2^f/f^* (which we term control or ‘wild-type’) and *Phd2^f/f^;THCre* (*Phd2ko*) animals with TH (tyrosine hydroxylase) restricted inactivation of *Phd2* in the carotid body and adrenal medulla and performed transcriptomic studies on those tissues. Pooled mRNA from 10 carotid bodies or adrenal medullas of animals of each genotype was analysed using RNA Seq.

Principal component analysis of this data indicated that differences between the carotid body and adrenal medulla are the largest source of variance between the datasets followed by the status of *Phd2* gene (Figure 1A). Although some established HIF target genes such as *Ldha* and *Bnip3* (Semenza et al. 1994; Bruick 2000) were upregulated in both the carotid body and adrenal medulla upon *Phd2* inactivation, overall changes in mRNA abundance by *Phd2* inactivation showed limited correlation (Spearman’s ρ∼0.1) between these tissues (Supplementary Figure 1A). Principal component analyses also indicated that *Phd2* inactivation in the adrenal medulla results in a modest overall shift in gene expression towards that of the carotid body (Figure 1A). To investigate this further, we focused on the subset of genes that were differentially expressed following *Phd2* inactivation in the adrenal medulla. Remarkably, our analysis showed that a large majority of the genes induced by *Phd2ko* in the adrenal medulla were more highly expressed in the carotid body compared to those in the adrenal medulla, whereas a large majority of those repressed by *Phd2ko* in the adrenal medulla were expressed at lower levels in the carotid body (Figure 1B, C). Since the same dataset for wild-type adrenal medulla is used in both comparisons (*Phd2ko* versus wild-type adrenal medulla, and carotid body versus adrenal medulla from wild-type mice), we went on to check whether this relationship is observed when using an independently generated comparator dataset comparing the carotid body versus adrenal medulla (Chang et al. 2015). The results were concordant (Supplementary Figure 1B versus Figure 1C), indicating the robustness of the finding. Overall, transcriptomic analyses demonstrate that *Phd2ko* in the adrenal medulla alters the expression of a subset of its transcripts toward a pattern that is similar to that in the carotid body. Notably, these include a number of genes involved in mitochondrial energy metabolism (*Cox4i2* and *Ndufa4l2*), G-protein signalling (*Rgs5* and *Adora2a*) and *Epas1* itself – all genes that are, or are implicated to be, involved in oxygen chemosensitivity (Hodson et al. 2016; Moreno-Dominguez et al. 2020; Pan et al. 2022)(see Supplementary Table 1 for full transcriptomic dataset and Supplementary Table 2 for a selection of genes that are, or have been implicated to be, involved in oxygen chemosensitivity).

**Figure 1.**
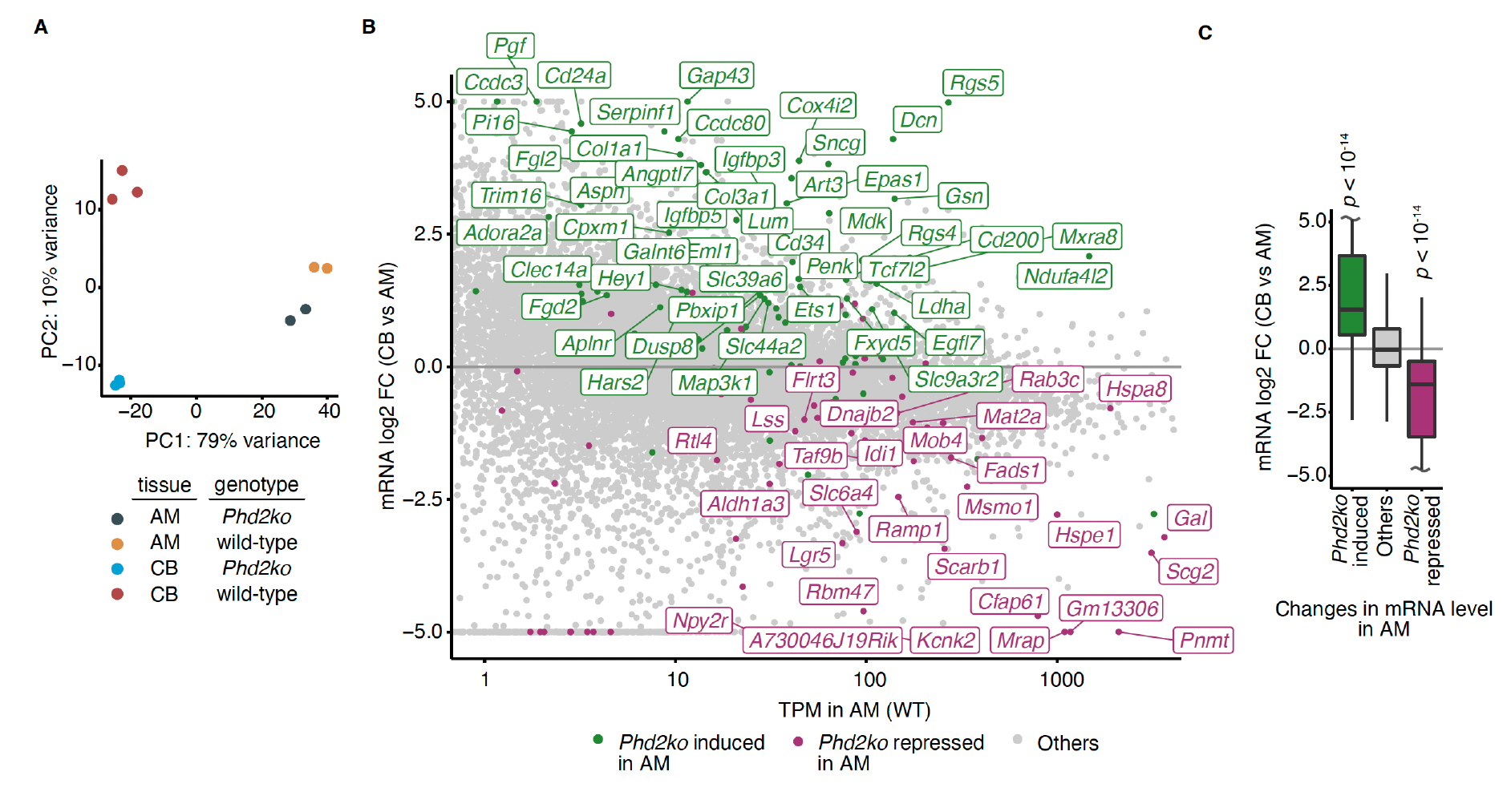
RNA Sequencing of *Phd2ko* (*Phd2^f/f^;THCre*) versus wild-type (*Phd2^f/f^*) adrenal medullas (AMs) and carotid bodies (CBs). (**A**) Principal Component Analysis (PCA) of bulk RNA Seq of CBs and AMs from *Phd2ko* and wild-type mice; RNA was extracted from 5 pairs of animals per biological replicate; n = 3 or 2 biological replicates for CBs or AMs, respectively. (**B**) Individual genes which are induced (green) or repressed (purple) by *Phd2ko* in the AM (the absolute fold change > log_2_(1.5) and FDR < 0.1, the likelihood ratio test) are overlaid onto genes differentially expressed in wild-type CB versus wild-type AM. Gene names are shown for a subset whose mRNA abundance in transcript per million (TPM) in wild-type or *Phd2ko* AM is greater than 10. (**C**) Data from (B) shown as a box plot. The fold difference in mRNA abundance in wild-type CB versus wild-type AM for genes induced (green) or repressed (purple) by *Phd2ko* in the AM was compared against that for all other genes (grey) using the two-sided Mann-Whitney U test. n = 108, 75 and 15,032, respectively.

### Phd2 inactivation confers oxygen chemosensitivity on chromaffin cells

In adult mammals the oxygen chemosensitivity associated with the carotid body is normally considered to be tightly restricted to the Type I cells in that organ. Other cells, including pulmonary vascular smooth muscle cells and neuroepithelial bodies, also manifest responses to hypoxia that occur on a similarly rapid time scale and may share some mechanisms (Youngson et al. 1993; Weir et al. 2005; Sommer et al. 2016). Whilst the adult adrenal medulla responds to neurogenic stimuli, it is not intrinsically responsive to hypoxia (Thompson et al. 1997). However, the above alterations in gene expression raised the interesting possibility that this paradigm might be broken by *Phd2* inactivation and confer carotid body-like chemosensitivity onto the adrenal medulla. Since hypoxia is established to promote Ca^2+^ influx in carotid body Type I cells following membrane depolarisation (Buckler 2015), intracellular Ca^2+^ concentration was recorded in response to superfusion of tissues with hypoxic solutions (equilibrated to 10-0% oxygen) or with a solution containing 45 mM K^+^ to maximally depolarise cells. Intracellular Ca^2+^ concentration ([Ca^2+^]_i_) fluctuations were detected by imaging tissues from transgenic mice expressing a genetically encoded calcium indicator: GCaMP6f recombined into the Rosa26 locus and preceded by a restricting ‘*lox-stop-lox’* sequence (known as *Ai95^f/+^*). These mice were intercrossed with THCre animals to generate *Ai95^f/+^;THCre* mice in which GCaMP6f expression is restricted to TH^+^ cells including those in the carotid body and adrenal medulla. We first confirmed that oxygen chemosensitivity is observed in carotid bodies from *Ai95^f/+^;THCre* mice. In these experiments, superfused whole carotid bodies were exposed to graded hypoxia by switching between superfusate equilibrated to 10, 5, 1, 0% oxygen, or a sham control in which a switch was made between perfusates with identical oxygenation. In these and all other experiments, superfusion with high (45 mM) K^+^ was included in the experimental protocol to define the regions that might respond to hypoxia and provide normalisation for other interventions (Supplementary Figure 2). Hypoxia resulted in a rapid (within seconds) and reversible increase in [Ca^2+^]_i_. Further, carotid body Type I cells responded to graded hypoxic stimuli with graded rises in [Ca^2+^]_i_, with responses to anoxia reaching ∼80% of the maximum increase in [Ca^2+^]_i_ achieved with high K^+^.

Next, we sought to test sensitivity to hypoxia in the adrenal medulla. To obtain a more consistent representation of the tissue (which is larger than, and has a different anatomy to, the carotid body, being encircled by the adrenal cortex), adrenal gland slices rather than whole tissues were used. Exposure to high K^+^ to induce membrane depolarisation resulted in a large increase in [Ca^2+^]_i_ which was used to define the region of interest for potential oxygen chemosensitivity (see Figure 2A). Unlike the carotid body, adrenal medullas from *Ai95^f/+^;THCre* mice that are wild-type for *Phd2* (referred to as ‘wild-type’ hereafter) were almost completely unresponsive to hypoxia or anoxia (Figure 2B, C). In striking contrast, adrenal medullas from *Phd2^f/f^;Ai95^f/+^;THCre* animals (referred to as ‘*Phd2ko*’ hereafter) manifest robust oxygen chemosensitivity when tested under identical conditions to adrenal medullas from wild-type mice (Figure 2B-D). The increase in [Ca^2+^]_i_ with hypoxia was rapid (within seconds) as well as reversible. Furthermore, the response was graded in response to the severity of the hypoxic challenge, as observed in the carotid body, with stimulation by anoxia reaching ∼35% of the maximum response obtained with high K^+^ (Figure 2B). These experiments therefore reveal that TH-restricted inactivation of *Phd2* is associated with the upregulation of a number of genes in the adrenal medulla whose expression is high in the carotid body and with the acquisition of oxygen-sensitive excitability, as manifest by rapid and reversible rises in [Ca^2+^]_i_ in response to hypoxia.

**Figure 2.**
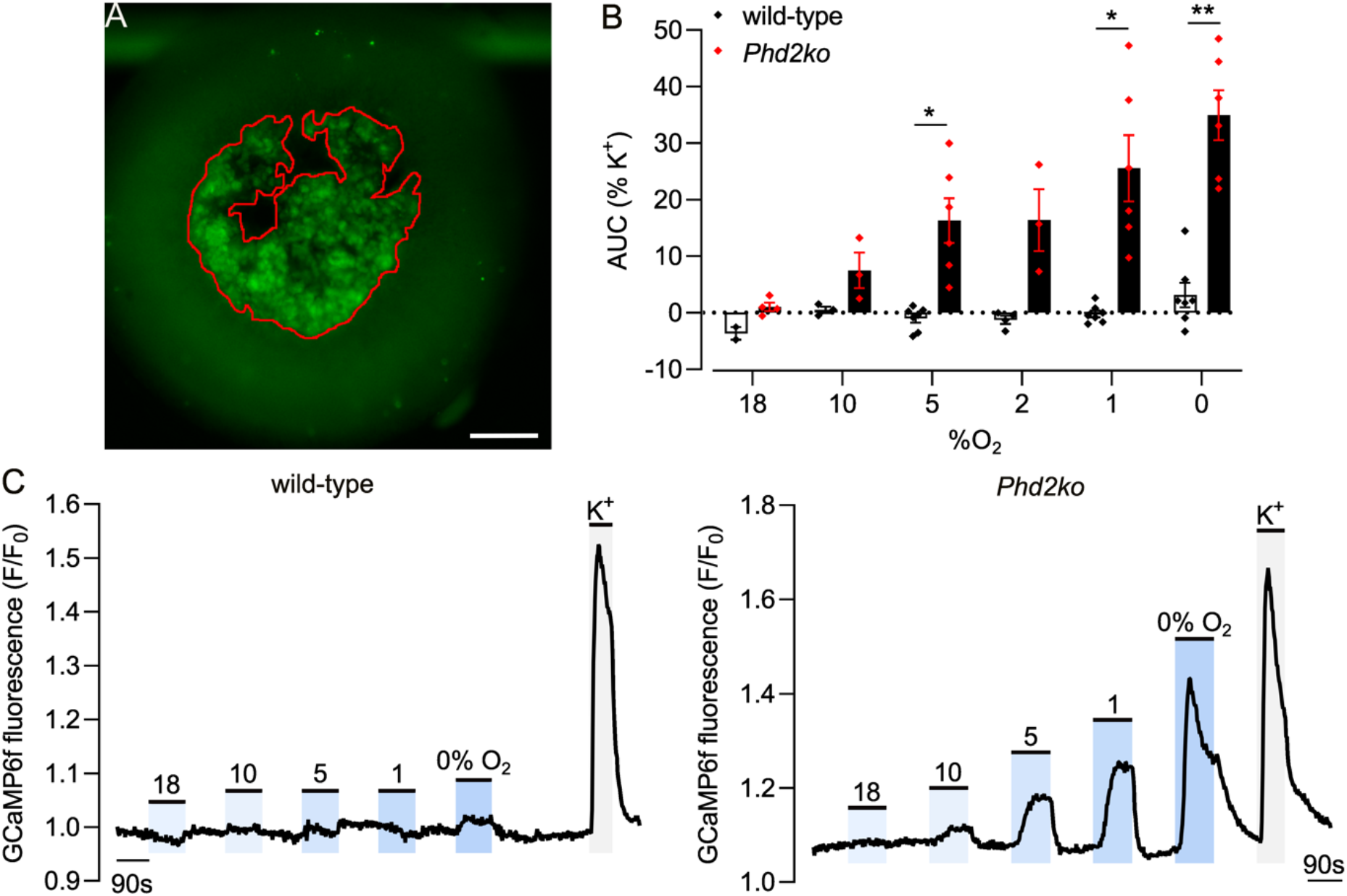
Ca^2+^ imaging showing oxygen chemosensitivity in *Phd2ko* (*Phd2^f/f^;Ai95^f/+^;THCre*), but not wild-type (*Ai95^f/+^;THCre*), adrenal medullas (AMs). (**A**) Representative image showing GFP fluorescence in an AM slice from a mouse expressing genetically encoded calcium indicator (GCaMP6f), the expression of which is restricted to TH^+^ cells (*Ai95^f/+^;THCre*, hereafter referred to as ‘wild-type’ in Figures 4, 5, 6 and 7) and perfused with 45 mM K^+^. Red outline shows an example of the K^+^-responsive region of interest from which fluorescence is quantified. Scale bar is 0.2 mm. (**B**) Average area under curve (AUC) corresponding to each hypoxic (or sham, 18% O_2_) stimulus in (C). Figures are normalised as % of AUC under 45 mM K^+^. Bars show mean ± S.E.M. with individual data points overlaid in this and subsequent figures. Data were analysed by a mixed-effects two-way ANOVA, with 18% O_2_ excluded from analysis: variation due to change in oxygen tension *P* < 0.0001, variation due to *Phd2* inactivation *P* = 0.0004; followed by a Sidak’s multiple comparisons test on pairwise comparisons at each oxygen level, * *P* < 0.05, ** *P* < 0.01. (**C**) Representative traces showing fluorescence (F) in the adrenal medulla (averaged across the K^+^-responsive region as per red outline in image (A)), background corrected and normalised to the fluorescence at the beginning of the recording (F_0_) to give F/F_0_; shaded areas highlight the time for which the indicated stimuli are applied: hypoxia (18-0% O_2_) or 45 mM K^+^ (in this and all subsequent figures depicting GCaMP6f recordings).

### Phd2 inactivation results in the formation of oxygen chemosensitive abdominal PGLs

In a previous study, we noted that *Phd2ko* results in the presence of collections of ectopic TH^+^ chromaffin cells, both within and surrounding the adrenal cortex (Eckardt et al. 2021). In light of the above findings, we sought to investigate this further. To that end, a systematic examination of the abdominal region was performed by transversely sectioning the area encompassing the superior to inferior mesenteric arteries and immunostaining for the presence of ectopic TH^+^ cells. Since the abdominal region contains several TH^+^ sympathetic ganglia, immunostaining for chromogranin A (CgA) was also performed to identify chromaffin cells. This revealed a large TH^+^ and CgA^+^ structure adjacent to the abdominal aorta and close to the inferior mesenteric artery (Figure 3A). This structure was absent in wild-type mice and its position and CgA positivity were strongly suggestive of a retained organ of Zuckerkandl (OZ). The OZ is a foetal structure that acts as the main source of catecholamines during development but disappears postnatally in wild-type mice (West et al. 1953; Coupland 1965; Schober et al. 2013). Our findings reveal that *Phd2ko* results in the apparent retention of an OZ-like CgA^+^ and TH^+^ structure in adult mice that was significantly enlarged compared to the OZ in either wild-type or *Phd2ko* foetal or newborn mice (Figure 3B and Supplementary Figure 3). Many of the TH^+^ and CgA^+^ cells had a ‘cleared’ appearance, previously described in paraganglioma (PGL)-like carotid bodies from these mice (Fielding et al. 2018) as well as OZ PGLs in humans (Subramanian and Maker 2006; Dossett et al. 2007). To pursue this resemblance to the carotid body, we tested the expression of selected genes that were upregulated in *Phd2ko* adrenal medulla, highly expressed in the carotid body and/or directly implicated in oxygen chemosensitivity (see above): *Epas1,* encoding HIF-2α; *Rgs5,* encoding a regulator of G-protein signalling; and *Cox4i2,* encoding an alternative regulatory subunit of mitochondrial cytochrome *c* oxidase (Figure 1B). All these genes were strongly expressed in this abdominal OZ PGL-like structure (Figure 3C).

**Figure 3.**
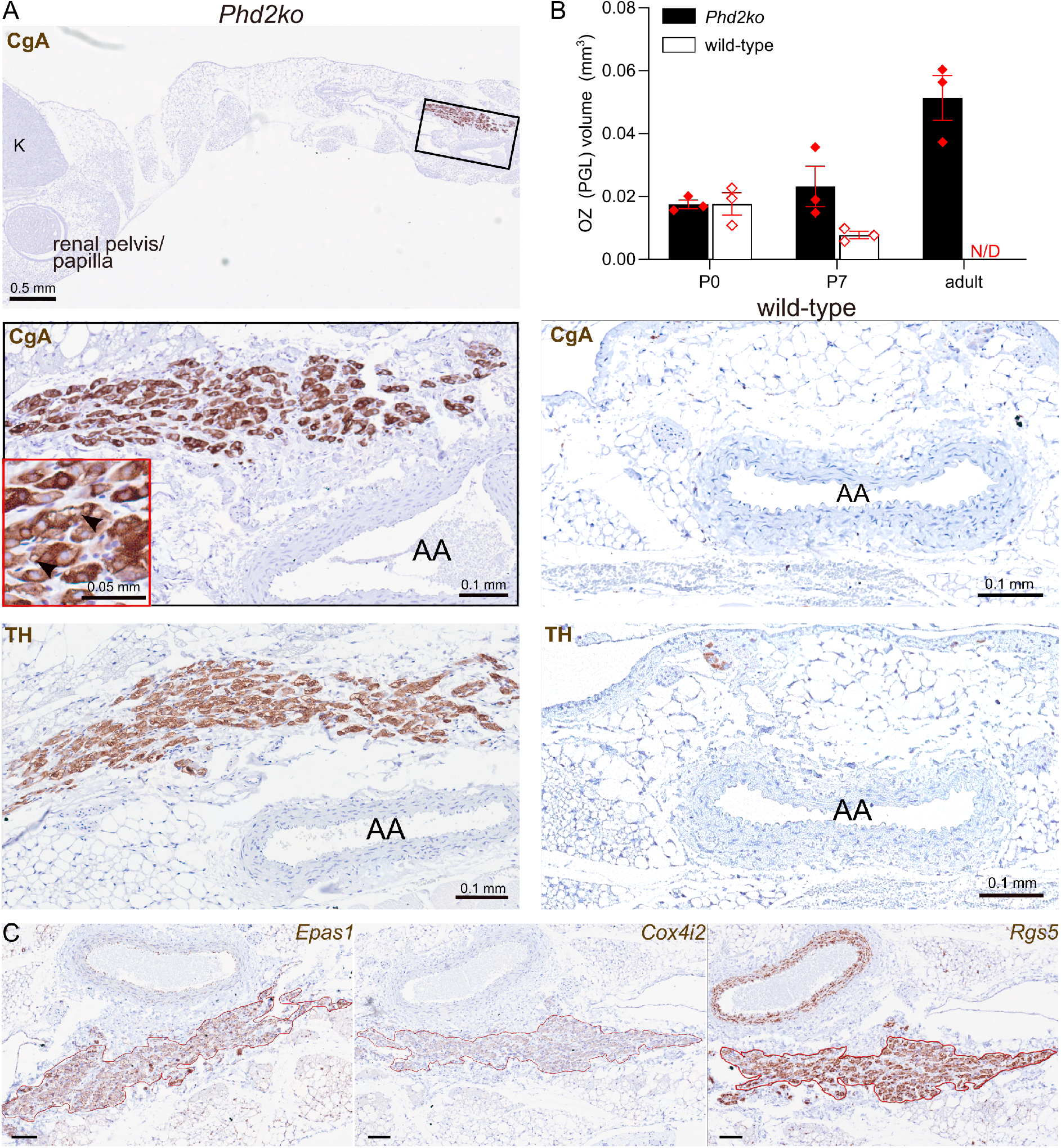
Abdominal, organ of Zuckerkandl paraganglioma (OZ PGL) in *Phd2ko* (*Phd2^f/f^;THCre*) but not wild-type (*Phd2^f/f^*) mice. (**A**) Chromogranin A (CgA, top left and middle two panels) and tyrosine hydroxylase (TH, bottom panels) immunohistochemistry in transverse sections of abdominal aorta (AA) and kidneys (K) in *Phd2ko* (left) and wild-type (right) adult mice. Low magnification image shows a transverse section of abdominal cavity, with the aorta-adjacent OZ PGL shown at higher magnification (see red insert for detailed cellular morphology showing ‘clearing’ within some CgA^+^ cells within the PGL). This structure is absent in a comparable region of the wild-type mouse. (**B**) OZ or OZ PGL volume based on CgA^+^ structure in abdominal cavities in neonatal (P0 and P7) and adult, *Phd2ko* and wild-type mice. No OZ PGL-like structure was detected in adult wild-type mice (N/D). (**C**) *In situ* hybridisation for *Epas1, Cox4i2* and *Rgs5* mRNA in the OZ PGL from an adult *Phd2ko* mouse. Scale bars are 0.1 mm.

We therefore sought to test whether the structure manifests oxygen chemosensitivity. Abdominal OZ PGLs were isolated from GCaMP6f expressing *Phd2^f/f^;Ai95^f/+^;THCre* (*Phd2ko*) mice and studied by perfusion of whole, subdissected OZ PGL preparations with hypoxic buffers. These experiments revealed a similar pattern of oxygen chemosensitivity to that observed in adrenal medullas from mice of the same genotype (Figure 4). Signals were again graded in accordance with oxygen tension, with stimulation by anoxia reaching a similar proportion of the response to high K^+^ (∼50%) to that observed in the adrenal medulla (Figure 4C).

**Figure 4.**
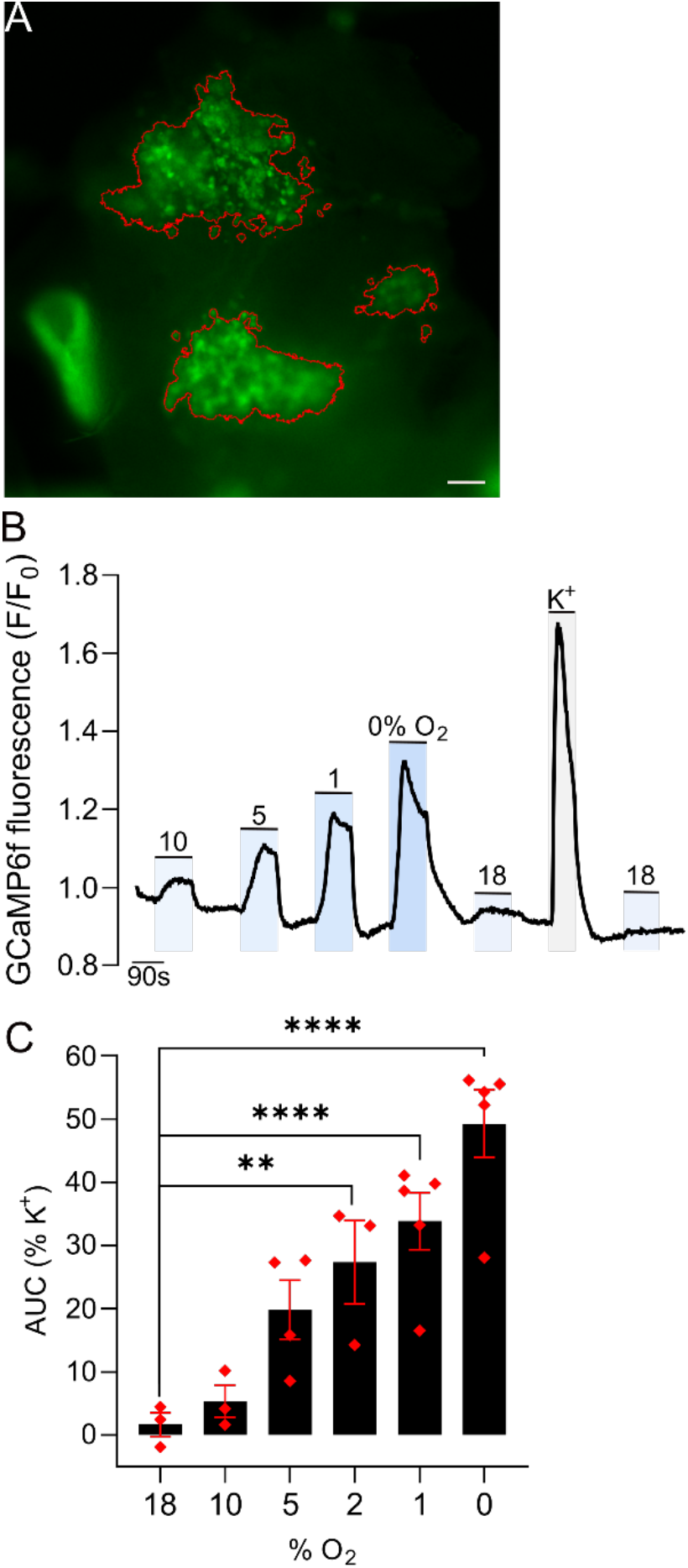
Oxygen chemosensitivity in the abdominal, organ of Zuckerkandl paraganglioma (OZ PGL) of a *Phd2ko* (*Phd2^f/f^;THCre*) mouse. (**A**) Representative image showing GFP fluorescence in an OZ PGL from an adult *Phd2ko* mouse perfused with 45 mM K^+^. Red outlines show the K^+^ responsive regions of interest, from which GCaMP6f fluorescence is quantified in (B) and (C). Scale bar represents 0.2 mm. Large bright-green structure visible in the bottom left corner is an adjacent blood vessel. (**B**) Representative trace showing F/F_0_ fluorescence in the K^+^ responsive regions as the tissue is exposed to the indicated stimuli: 10-0% O_2_, 45 mM K^+^ or sham (18% O_2_). (**C**) Average area under curve (AUC) normalised to that in response to 45 mM K^+^ in *Phd2ko* OZ PGLs. Data were analysed by a one-way, repeated measures ANOVA, variation due to oxygen tension *P* = 0.0035, followed by Dunnett’s multiple comparisons test comparing each stimulus to 18% O_2_, ** *P* < 0.01, **** *P* < 0.0001.

### Doxapram and CO_2_ sensitivity of Phd2ko AMs and abdominal OZ PGLs

Next, we sought to further characterise the chemosensitivity of these adrenal medullas and OZ PGL-like structures in *Phd2ko* mice, in particular their resemblance to carotid body chemosensitivity. We first tested responses to doxapram (1-ethyl-4-(2-morpholin-4-ylethyl)-3,3-diphenylpyrrolidin-2-one), a central respiratory stimulant, whose mode of action is thought to be via direct stimulation of carotid body Type I cells (O’Donohoe et al. 2018). Adrenal medullas were perfused with 50 μM doxapram, which is reported to inhibit TASK channel activity and evoke an increase in cytosolic Ca^2+^ in dissociated carotid body Type I cells (O’Donohoe et al. 2018). Second, we tested responses to acid/CO_2_, which is also known to excite the carotid body (Iturriaga and Lahiri 1991; Rocher et al. 1991; Buckler and Vaughan-Jones 1994). Adrenal medullas (and to a lesser extent the OZ PGLs) from *Phd2ko*, but not wild-type, mice were sensitive to both 50 μM doxapram and 10% CO_2_ (Figure 5).

**Figure 5.**
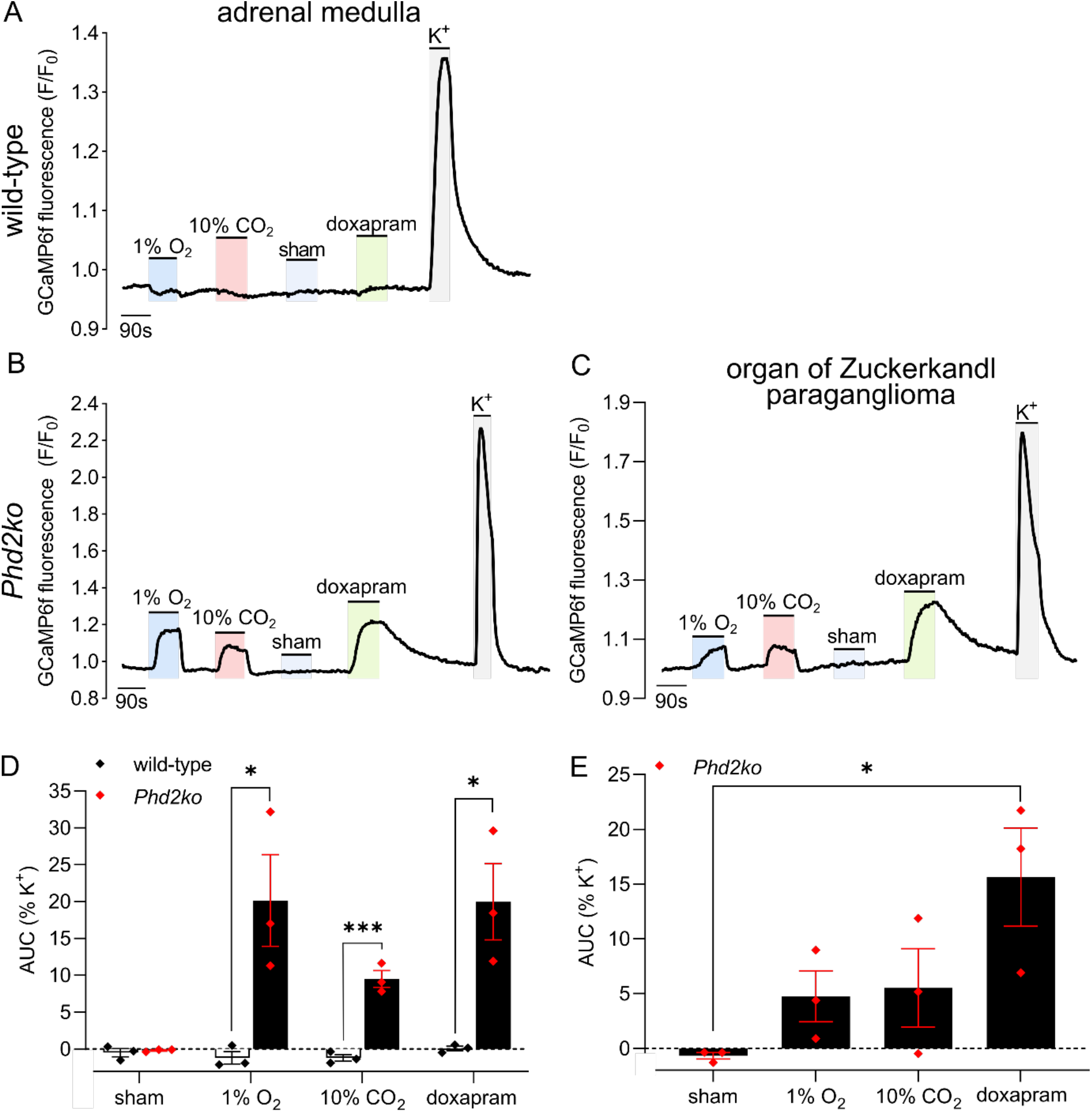
*Phd2* inactivation induces Ca^2+^ mobilisation in chromaffin cells in response to 10% CO_2_ and doxapram. Representative traces of GCaMP6f F/F_0_ fluorescence in the K^+^ responsive region of (**A**) wild-type (*Ai95^f/+^;THCre*) and (**B**) *Phd2ko* (*Phd2^f/f^;Ai95^f/+^;THCre*) adrenal medulla (AM) and (**C**) *Phd2ko* organ of Zuckerkandl paraganglioma (OZ PGL), as the tissues are exposed to indicated stimuli: 1% O_2_, 10% CO_2_, sham test solution (18% O_2_), 50 μM doxapram or 45 mM K^+^. (**D, E**) Average area under curve (AUC) normalised as % of AUC with 45 mM K^+^ in (D) wild-type and *Phd2ko* AM slices and (E) OZ PGL. Data were analysed by (D) two-tailed, unpaired Student’s *t-*tests, * *P* < 0.05, *** *P* < 0.001; (E) one-way ANOVA, variation due to oxygen tension *P* = 0.0334, followed by Dunnett’s multiple comparisons test against the sham*, * P* < 0.05.

### Activation of HIF-2α is sufficient to induce oxygen chemosensitivity

Inactivation of the principal oxygen sensor PHD2 leads to stabilisation of HIF-α subunits and activation of HIF. Although the mechanisms are unclear, several pieces of data have implicated one isoform: HIF-2α, as being necessary for carotid body development and physiological function, and for the paraganglioma-like phenotypes associated with inactivation of *Phd2* (Hodson et al. 2016; Fielding et al. 2018; Macias et al. 2018; Cheng et al. 2020; Moreno-Dominguez et al. 2020; Eckardt et al. 2021). Together with the above observations on induction of oxygen chemosensitivity, these findings led us to consider whether overexpression of HIF-2α alone might be sufficient to induce oxygen chemosensitivity in the adrenal medulla. To address this, a gain-of-function *Hif-2α* allele (*Hif-2αdPA*), in which the two sites of prolyl hydroxylation in HIF-2α are substituted with alanine residues to prevent hydroxylation and hence convey resistance to degradation by the VHL ubiquitin-proteasomal pathway, was used (Kim et al. 2006). Activation of this transgene in *Hif-2αdPA^f/+^*;*TH*Cre mice (referred to as ‘*Hif-2αdPA*’ hereafter) leads to HIF-2α overexpression in TH^+^ cells of the carotid body and adrenal medulla, which was confirmed in the adrenal medulla, carotid body and organ of Zuckerkandl by immunohistochemistry for HIF-2α as well as by immunoblotting for the hemagglutinin (HA) tag present on HIF-2αdPA (Supplementary Figures 4, 5). To demonstrate the activity of this transgene, we first tested for effects on the carotid body. Consistent with earlier work (Macias et al. 2014), these experiments demonstrated that *Hif-2αdPA* mice developed enlarged paraganglioma-like carotid bodies (Supplementary Figure 6A, B). In line with this we also observed an increase in ventilatory sensitivity to hypoxia (Supplementary Figure 6C, D).

Importantly, anatomical and histological examination of the adrenal glands, peri-adrenal and abdominal regions of these animals also revealed abnormalities that were essentially identical to those observed in the *Phd2ko* mice. Particularly striking was the retention of OZ PGL-like tissues in the adult (Figure 6A, B). As with *Phd2ko* animals, this tissue expressed increased levels of *Rgs5*, *Cox4i2*, as well as *Epas1* mRNA (Supplementary Figure 7). In the adrenal medulla, a switch in the cellular pattern of gene expression towards apparently immature phenylethanolamine N-methyltransferase (*Pnmt)^-^/Rgs5^+^/Epas1^+^* cells was observed in HIF-2α overexpressing cells (Figure 7A), as is also observed in *Phd2ko* animals (where it is reversed by inactivation of *Hif-2α*, but not *Hif-1α*)(Eckardt et al. 2021). Robust induction of oxygen chemosensitivity closely similar to that observed in *Phd2ko* mice was observed both in the adrenal medullas and the abdominal OZ PGLs of *Hif-2αdPA* (*Hif-2αdPA^f/+^;Ai95^f/+^;THCre*) animals (Figures 6 and 7). Thus, stabilised HIF-2α is sufficient to drive a paraganglioma-like oxygen chemosensitive phenotype in chromaffin cells which, under the conditions of these experiments, was essentially identical to that observed with *Phd2* inactivation.

**Figure 6.**
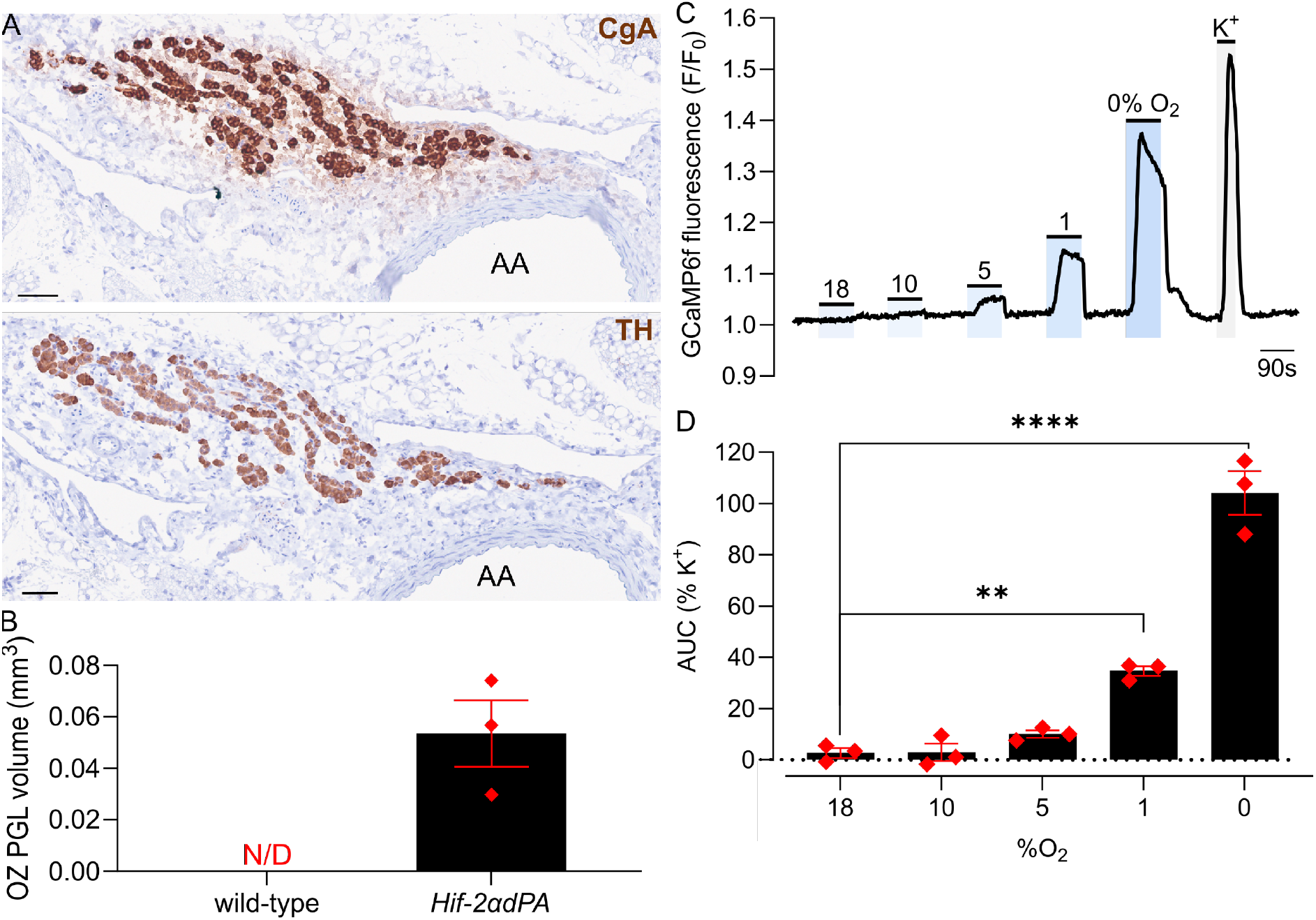
Abdominal, organ of Zuckerkandl paraganglioma (OZ PGL) in *Hif-2αdPA* (*Hif-2αdPA^f/+^;THCre*) mice. (**A**) Chromogranin A (CgA, top panel) and tyrosine hydroxylase (TH, bottom panel) immunohistochemistry in OZ PGL shown in transverse sections of abdominal aorta (AA) in adult *Hif-2αdPA* mice. Scale bars are 0.05 mm. (**B**) OZ PGL volume based on CgA^+^ structures in abdominal cavities in adult *Hif-2αdPA* mice. No OZ PGL-like structure was detected in adult wild-type mice (N/D). (**C**) Representative trace showing F/F_0_ fluorescence in the K^+^ responsive regions, as the tissue is exposed to indicated stimuli: hypoxia (18-0% O_2_) or 45 mM K^+^. (**D**) Average area under curve (AUC) normalised as % of AUC under 45 mM K^+^ in *Hif-2αdPA* OZ PGLs (with the sham control used for 18% O_2_). Data were analysed by a one-way ANOVA, variation due to oxygen tension *P* < 0.0001, followed by a Dunnett’s multiple comparisons test against 18% O_2_, ** *P* < 0.01, **** *P <* 0.0001.

**Figure 7.**
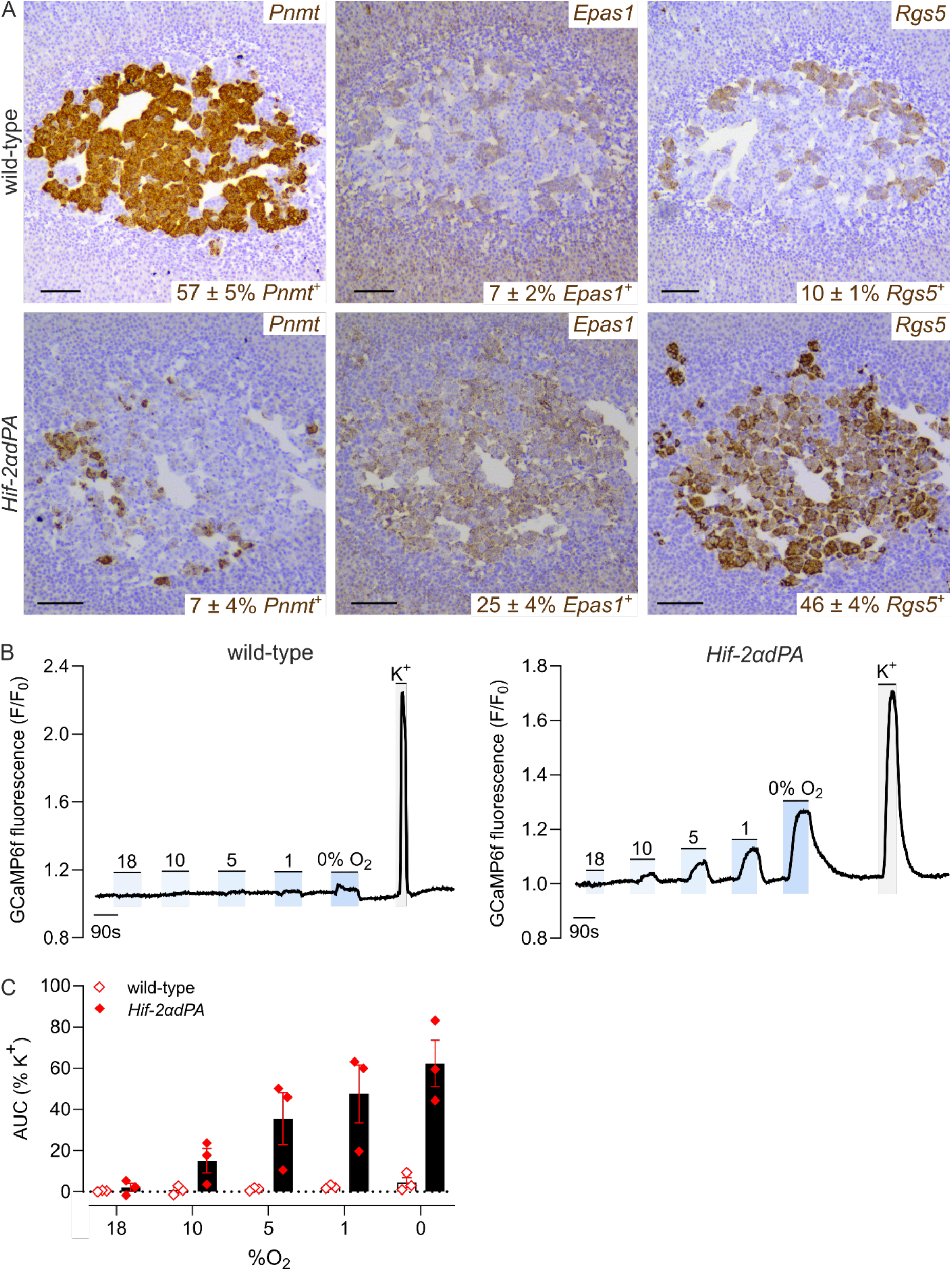
Gene expression and oxygen chemosensitivity in wild-type and *Hif-2αdPA* (*Hif-2αdPA^f/+^;Ai95^f/+^;THCre*) adrenal medullas (AMs). (**A**) *In situ* hybridisation for *Pnmt*, *Epas1* (*Hif-2α*) and *Rgs5* (brown) in adjacent sections from wild-type and *Hif-2αdPA* mice. Harris haematoxylin counterstain (purple). Scale bars are 0.1 mm. Proportion of AM area expressing each mRNA is displayed at the bottom as mean ± S.E.M., n = 3. For each mRNA, mean area of expression was compared between genotypes using an unpaired Student’s *t*-test, *Pnmt P* = 0.0015, *Epas1 P* = 0.0138, *Rgs5 P* = 0.0008. (**B**) Representative traces showing F/F_0_ fluorescence in wild-type versus *Hif-2αdPA* AMs as the tissue is exposed to the indicated stimuli: 10-0% O_2_, 45 mM K^+^ or sham (18% O_2_). (**C**) Average area under curve (AUC) to each hypoxic (or sham, 18% O_2_) stimulus (as recorded in (B)) normalised as % of AUC under 45 mM K^+^. Data were analysed by a two-way ANOVA with 18% O_2_ excluded from analysis: variation due to change in oxygen tension *P* = 0.0015, variation due to HIF-2α activation *P* = 0.0238, variation due to interaction between change in oxygen tension and HIF-2α activation *P* = 0.0003; followed by a Sidak’s multiple comparisons test on pairwise comparisons at each oxygen level.

## Discussion

Our findings establish that activation of hypoxia signalling pathways, either by inactivation of the regulatory HIF prolyl hydroxylase PHD2 or by overexpression of one of its targets, HIF-2α, is sufficient to induce oxygen chemosensitivity in the adult adrenal medulla. We observed that this was associated with a bidirectional shift in expression of selected genes within the adrenal medulla that coincides with differences in gene expression between the normal carotid body and the normal adrenal medulla.

Interestingly, this pattern of gene expression included genes with proposed functions in generating or tuning hypoxic signals from the mitochondria. Thus, expression of *Cox4i2*, an alternative isoform of *Cox4*, which is a nuclear encoded regulatory subunit of the cytochrome *c* oxidase complex and *Ndufa4l2*, another nuclear encoded component of this complex, were both increased in the oxygen chemosensitive adrenal medullas of *Phd2ko* mice (Supplementary Table 2). Our findings therefore strengthen the hypothesis that these genes, and their actions on the kinetic properties of cytochrome *c* oxidase, are important determinants of oxygen chemosensitivity (Moreno-Dominguez et al. 2020). Another intriguing finding was the striking upregulation of *Rgs5* transcripts, which encode a regulator of G-protein signalling that is a target of the recently recognised oxygen sensitive N-terminal cysteine dioxygenase ADO (2-aminoethanethiol dioxygenase)(Masson et al. 2019).

Remarkably, the great majority of transcripts that were dysregulated by *Phd2ko* in the adrenal medulla were similarly up or down-regulated when the normal carotid body was compared with the normal adrenal medulla. The exact relationship of these changes in gene expression (provided as a resource in Supplementary Table 1) to the induced oxygen chemosensitivity in *Phd2ko* adrenal medullas will require further study. Particularly significant in the current study, however, was the sufficiency of forced expression of HIF-2α to create oxygen chemosensitivity. HIF-2α was identified as one of the transcripts whose expression is increased in the *Phd2ko* adrenal medulla. It is expressed at strikingly high levels in the carotid body (Tian et al. 1998; Zhou et al. 2016; Fielding et al. 2018) and it is necessary for the development and oxygen chemosensitivity of that organ and for the enhanced hypoxic ventilatory responses mediated by the carotid body in hypoxic acclimatisation (Hodson et al. 2016; Fielding et al. 2018; Macias et al. 2018; Cheng et al. 2020; Moreno-Dominguez et al. 2020). Taken together with these observations, the current findings suggest that, amongst its other functions, HIF-2α appears to be integral to the specification of oxygen chemosensitivity in sympathoadrenal or chromaffin cells.

Interestingly, the induced chemosensitivity in *Phd2ko* and *Hif-2α* overexpressing adrenal medullas, and that of the retained OZ-like structures, was not limited to oxygen chemosensitivity, but included responsiveness to high CO_2_ and doxapram. Since equilibration of the bicarbonate buffer with high CO_2_ will reduce pH, we are not able to distinguish whether the direct sensitivity is to acid or CO_2_ itself. Nevertheless, both high acid/CO_2_ and doxapram are established stimulants of the Type I chemosensitive cells in the carotid body, where the targets are considered to be the TASK (Twik-related acid-sensitive K^+^) channels, TASK1 and TASK3 (Cotten et al. 2006; O’Donohoe et al. 2018). However, TASK1 (*Kcnk3*) and TASK3 (*Kcnk9*) transcripts were neither overexpressed in the carotid body versus adrenal medulla nor induced in *Phd2ko* adrenal medullas (Supplementary Table 2).

Several pieces of information suggest that the alterations we have observed following intervention on the PHD2/HIF-2 pathway reflect reversion to a foetal-like state. First, the alteration in gene expression involves loss of adrenal medullary expression of *Pnmt*, which has been considered to be a marker of chromaffin cell differentiation (Verhofstad et al. 1979). Second, the changes involve retention of OZ-like structures that normally characterise the foetal state. Third, the physiological characteristics, chemosensitivity to oxygen and CO_2_, reflect those that have been reported for foetal adrenal tissues (Thompson et al. 1997).

Though further work will be required to understand the exact basis of this phenotypic switch, the current findings have biochemical, physiological and clinical implications. By recording the induction of oxygen chemosensitivity in a previously unresponsive tissue the work enhances the power of association studies between gene expression and phenotype. More generally, the work provides a new model for the study of oxygen chemosensitivity, including future studies as to which of the downstream targets of HIF-2α are required for oxygen chemosensitivity. It also has potential implications for the phenotype of ‘pseudohypoxic’ pheochromocytoma, which typically manifests upregulation of genes in the HIF pathway and is associated with mutations in *HIF-2α*, *PHD2*, *PHD1*, *VHL* and specific subunits of tricarboxylic acid cycle enzymes, succinate dehydrogenase *SDHx*, which are frequently germline or early post-zygotic, so potentially developmental in origin (Crona et al. 2017; Dariane et al. 2021). It will now be important to determine if oxygen chemosensitivity is a characteristic feature of these tumours, as awareness of factors stimulating catecholamine release from these tumours is critical to safe clinical management. Further, given the association between hypoxic activation of chemosensing and cell proliferation in the carotid body (Platero-Luengo et al. 2014), the work also raises an interesting question as to whether oxygen chemosensitivity contributes to the growth and development of these tumours.

In summary, our work reveals a new link between two key systems of oxygen sensing. It should be important in directing future work on the pathogenesis of paraganglioma/pheochromocytoma development and more generally in investigating the fundamental physiological challenge of maintaining oxygen homeostasis.

## Materials and methods

### Animals

Animal experimental protocols were approved by the University of Oxford Medical Science Division Ethical Review Committee and are compliant with the UK Home Office Animals (Scientific Procedures) Act 1986. Experiments were performed on ∼2 month old mice of the relevant genotype and sex-matched controls. Mice were kept in individually ventilated cages with free access to water and food. *Phd2^f/f^* and *THCre* alleles are as described (Lindeberg et al. 2004; Mazzone et al. 2009); *Ai95^f/+^* and *Hif-2αdPA^f/+^* were obtained from Jackson laboratories (strains: 028865, 009674) and are as described (Kim et al. 2006; Madisen et al. 2015). Each mouse line was backcrossed with C57BL/6 for at least five generations.

### Immunohistochemistry

Tissue processing and immunohistochemistry procedures were as described previously (Bishop et al. 2013; Hodson et al. 2016; Fielding et al. 2018). Animals were culled with overdose of isoflurane or cervical dislocation and tissues dissected and placed in 4% paraformaldehyde (w/v in phosphate buffered saline, PBS) and left to fix at room temperature overnight, then transferred into 70% ethanol and stored at 4°C until processing. Tissues were processed by gradual dehydration in 70%, 90% and 100% ethanol, then Histo-Clear II (National Diagnostics) and finally wax at 60°C in an automated tissue processor (Excelsior AS, Thermo Scientific), then embedded in paraffin and sectioned with a Microm HM 355S Microtome (Thermo Scientific) to 4 μm thickness. To detect catecholaminergic cells, paraffin sections were immunostained with anti-CgA antibody (1:500 ab68271, Abcam) using the Envision+ Kit (Dako) according to manufacturer’s instructions, and counterstained with modified Harris Hematoxylin (Epredia) differentiated with 1% acetic acid (Merck) for optimal stain intensity. Stained tissues were scanned with NanoZoomer S210 slide scanner (Hamamatsu) and OZ volume estimated by calculating CgA^+^ area using NDP.View2 software (Hamamatsu) as described (Fielding et al. 2018).

### In situ hybridisation

RNA was detected in formalin-fixed paraffin-embedded tissues (as prepared above) using the RNAScope manual assay (ACDBio) as per manufacturer’s instructions, with the following gene-specific probes: *Epas1* (catalogue number: 314371), *Rgs5* (430181), *Cox4i2* (497901). Harris Hematoxylin (Epredia), differentiated with 1% acetic acid for optimal intensity, was used as a counterstain.

### RNA extraction and RNA Sequencing

CBs and AMs were pooled from five animals per genotype (i.e. ten in total for each tissue) to obtain sufficient material and RNA was extracted using RNEasy Micro Plus kit (Qiagen) according to manufacturer’s instructions and as described previously (Gao et al. 2017). Tissues were sub-dissected into RNAProtect Tissue Reagent (Qiagen) and placed on ice for 2-5 h until all tissues were collected. 1 volume of PBS in Diethyl Pyrocarbonate-treated H_2_O was then added to Tissue Protect and tissues centrifuged at 16,100 g for 1 min to collect at the bottom of the tube, allowing removal of the Tissue Protect and PBS mix. Subsequently, 300 μl of ice-cold RLT Plus buffer with 20 μl of 2 M dithiothreitol (DTT) per 1 ml was added and tissues homogenised using the Pro200 homogeniser (ProScientific) in: 5 x 6 s pulses for CBs and 4 x 5 s pulses for AMs. Lysate was then centrifuged for 3 min at 16,100 g, 4°C and supernatant transferred into a clean, RNAse-free tube then immediately frozen on dry ice. Lysates were stored at −80°C for 1-7 days before continuing with RNA extraction. Lysates were thawed on ice and the remaining steps from Qiagen kit were carried out at room temperature. RNA was eluted with 14 μl RNAse-free H_2_O and quality assessed using the High Sensitivity RNA ScreenTape (Agilent), as per manufacturer’s instructions. Remaining RNA was used immediately or stored at −80°C. Ultra-low library preparation and RNA sequencing using the NovaSeq6000 platform were performed by the Genomics Facility at the Wellcome Centre for Human Genetics, University of Oxford.

### RNA sequencing data processing and reference data

Based on the RNA sequencing data, the transcript abundance was calculated using Salmon (1.9.0)(Patro et al. 2017) with the reference transcript data by GENCODE (v31)(Frankish et al. 2021). K^+^ channel related genes were defined as those associated with the GO term, GO:0005267 (K^+^ channel activity).

### RNA sequencing data analysis (principal component analysis)

The transcript count data of genes was transformed using the vst function of DESeq2 without considering the sample information, and the principal component analysis was performed using the plotPCA function of DESeq2 (Love et al. 2014).

### RNA sequencing data analysis (identification of differentially expressed genes)

The fold change and statistical significance of differential gene expression between two tissues or upon *Phd2*ko were calculated using DESeq2. For the analysis of the statistical significance of differential gene expression upon *Phd2ko*, the difference between batches of samples was considered using the likelihood ratio test whenever possible. Genes with the false discovery rate < 0.1 and the absolute fold change > log_2_(1.5) were defined as differentially expressed. Genes with a very low expression level were identified by DESeq2 and these genes were excluded from the analyses. The scripts and pipeline used for the analysis of RNA Seq are available on GitHub (https://github.com/YoichiroSugimoto/20230620_Phd2KO_in_AM_and_CB). Sequence data generated from this study are available from ArrayExpress (E-MTAB-13163).

### Carotid bifurcation and adrenal gland preparation for live Ca^2+^ imaging

Mice were culled by terminal anaesthesia (or cervical dislocation if CBs not used) and carotid bifurcations, aortas and adrenal glands were collected into ice-cold PBS. CBs and OZ PGLs were then sub-dissected in ice-cold PBS and placed on a nylon mesh (Warner Instruments) in F12 media (Gibco) with Foetal Bovine Serum (Gibco), Penicillin/Streptomycin and L-Glutamine, then kept at 37°C, 5% CO_2_ until imaging (for up to 4 h). Adrenal glands were embedded in 5% low-melting-point agarose (BP1360, Fisher Scientific) in modified Krebs-Henseleit buffer (25 mM NaHCO_3_, 117 mM NaCl, 1 mM NaH_2_PO_4,_ 11 mM D-Glucose, 4.5 mM KCl, 1 mM MgCl_2_, 25 mM HEPES) at 39°C and allowed to cool on ice for ∼5 min. 150 μm slices were obtained with a VT100S Vibratome (Leica) using Feather FA-10 carbon blades (Labtech) vibrating at 80 Hz, 0.2 mm/s. Slices were placed in DMEM media (Gibco) then kept at 37°C, 5% CO_2_ for at least 1 h before imaging (for up to 4 h).

### Fluorescent imaging

Tissues were imaged in a custom-built gravity-based perfusion system, with a wide-field Leica DMi8 microscope (Leica Microsystems CMS), configured with a pE340^fura^ light source (CoolLED), a GFP filter cube (excitation filter 470 ± 40 nm, 495 nm dichroic mirror, emission filter 525 ± 50 nm) and a monochrome Prime 95B camera (Photometrics). Krebs-Henseleit buffer (25 mM NaHCO_3_, 117 mM NaCl, 1 mM NaH_2_PO_4,_ 11 mM D-Glucose, 4.5 mM KCl, 1 mM MgCl_2_)-based test solutions were bubbled with gases using 3-channel Gas Blender 100 Series mixers (100 series, MCQ Instruments) for at least 2 h prior to the start of the experiment; CaCl_2_ was added after ∼2 h of bubbling with 5% CO_2_ to a final concentration of 1 mM to prevent calcium carbonate precipitation. Solutions were heated in-line upstream of the tissue chamber to achieve a temperature of ∼37°C in the chamber. Tissues were placed in the perfusion chamber and immobilised with a slice anchor (SHD-22L/15 or SHD-22L/10, Warner Instruments) and control solution was supplied until the start of the experiment. GFP images were obtained every 2 s with a x40 oil-immersive objective (HC PL FLUOTAR 40x/1.30 OIL 340 nm, Leica Microsystems CMS) used for CBs and a x10 objective (N PLAN 10x/0.25 achromatic corrected objective, Leica Microsystems CMS) used for OZ and AM. Bright-field images of the AM were taken prior to the start of recording to select the AM area for analysis. Baseline fluorescence was measured for at least 60 s prior to the application of the first test. Test solutions were applied for 90 s (or up to 60 s for K^+^), with at least 120 s rest in control solution between tests. For composition of test solutions, see Supplementary Table 3. Raw data were analysed using Fiji software (NIH). Average GFP fluorescence intensity across the region of interest was quantified across the z-stack and plotted against time. For background correction, small regions on-tissue but away from K^+^-responsive cells were selected.

### Statistical analysis

All statistical analyses were carried out in Graphpad Prism 9.0 or R (4.2.1) software. Details of statistical tests are included in the figure legends. All data are presented as mean ± S.E.M..

## Acknowledgements

Funding for the work was received from the Oxford Branch of Ludwig Cancer Research, the Wellcome Trust (106241/Z/14/Z) and the Paradifference Foundation. This work was also supported by the Francis Crick Institute, which receives its core funding from Cancer Research UK (FC001501), the UK Medical Research Council (FC001501), and the Wellcome Trust (FC001501).

## Author contributions

Experiments were designed by MPB, HAJ, YS, KJB, PJR, TPK and TB. Data were collected and analysed by all authors. Manuscript was prepared by MPB, HAJ, YS, PJR, TPK and TB and reviewed by all authors. Figures were prepared and statistical analyses performed by MPB, HAJ and YS with input from other authors. PJR, TPK and TB conceived the study and managed the project. MBP, HAJ and YS are co-first authors; PJR, TPK and TB are co-senior authors.

