## Supplementary Figures for "*Hif-2α* programmes oxygen chemosensitivity in chromaffin cells"

### Supplementary materials

Maria Prange-Barczynska[1,2], Holly A. Jones[1,2], Yoichiro Sugimoto[3,4,5], Xiaotong Cheng[1,2], Joanna D. C. C. Lima[1,2], Indrika Ratnayaka[2], Gillian Douglas[6], Keith J. Buckler[7], Peter J. Ratcliffe[1,2,3], Thomas P. Keeley[1] and Tammie Bishop[1,2]

[1]Target Discovery Institute, University of Oxford, Oxford, UK

[2]Ludwig Institute for Cancer Research, University of Oxford, Oxford, UK

[3]The Francis Crick Institute, London, UK

[4]Max Delbrück Center for Molecular Medicine, Berlin, Germany

[5]DZHK (German Centre for Cardiovascular Research), Partner Site Berlin, Berlin, Germany

[6]BHF Centre of Research Excellence, Division of Cardiovascular Medicine, Radcliffe Department of Medicine, John Radcliffe Hospital, University of Oxford, Oxford, UK

[7]Department of Physiology, Anatomy & Genetics, Parks Road, Oxford, UK

**Authorship note:** MPB, HAJ and YS are joint first authors. PJR, TPK and TB are co-senior authors.

;

**Supplementary methods**

Experiments presented in Supplementary materials were carried out according to the same protocols as described in the main manuscript, except for the following:

*HA immunoblotting*

Tissues were dissected, lysed in sodium dodecyl sulfate (SDS) lysis buffer (50mM Tris pH 6.8, 2% SDS, 10% glycerol) and sonicated to homogeneity. Samples were normalised for protein concentration using a BCA assay (Pierce, Thermo Fisher Scientific), then mixed with Laemmli sample buffer. Proteins were separated by SDS-polyacrylamide gel electrophoresis then transferred onto a polyvinylidene difluoride membrane (Immobilon-P, Millipore). This was blocked in 4% fat free milk (w/v in phosphate buffered saline, PBS, containing 0.1% Tween20) then incubated overnight with HRP-conjugated HA (3F10, Sigma Aldrich) diluted 1:500 in milk. Chemiluminescence substrate (West Dura, 34076, Thermo Fisher Scientific) was added and bound antibody was detected using a ChemiDoc XRS+ imaging system (BioRad). β-actin (ab49900, Abcam) was used as a loading control.

*Multi-layer indirect labelling for HIF-2α*

For the detection of HIF-2α, a new signal amplification system was developed in the laboratory. 4 µm formaldehyde-fixed paraffin-embedded sections were deparaffinised, rehydrated and subjected to heat-induced epitope retrieval with the Dako Target Retrieval Solution pH 6.0 (S2369, Agilent) for 30 min at 120°C. Sections were blocked for non-specific protein, avidin, and biotin binding and were incubated with anti-HIF-2α primary antibody (Wiesener et al. 2003)(PM8; 1:3000) overnight at 4°C. Indirect labelling was performed by serial incubations with secondary HRP-conjugated anti-rabbit antibody (Dako K4003, Agilent; 30 min room temperature, RT), tertiary biotin-conjugated anti-HRP antibody (Rockland 200-4638-0100; 1:250; 2 h RT) and quaternary HRP-conjugated Streptavidin (BD Pharmingen 551321; 1 h RT). Slides were developed using 3,3′-diaminobenzidine chromogen and counterstained with haematoxylin.

*Plethysmography*

Hypoxic ventilatory response (HVR) was assessed by whole-body plethysmography in awake animals (Bishop et al. 2013; Cheng et al. 2020): mice were placed in Buxco whole-body plethysmographs (PLY4211, 600 ml, DSI) and allowed to acclimatise to the new environment for 30 min with plethysmographs unsealed and filled with room air. 1.5 $L \cdot min^{-1}$ medical air flow was initiated ~3 min prior to the start of the recording and minute ventilation was measured for 5 min in air, followed by 5 min at 10% $O_2$/3% $CO_2$/balance $N_2$ (supplied from a pressured gas cylinder, BOC) and another 5 min at medical air. Data were normalised to animal's body weight recorded immediately prior to placing the animal in the plethysmograph and plotted against time. HVR was quantified as the difference between the average minute ventilation during the first minute of stable hypoxia (excluding the first 30s after switching the gas supply) and the average minute ventilation during the last minute of normoxia.

**Supplementary Tables**

**Supplementary Table 1. RNA Sequencing of *Phd2ko* (*Phd2$^{f/f}$;THCre*) versus wild-type (*Phd2$^{f/f}$*) adrenal medullas (AMs) and carotid bodies (CBs).** RNA expression in *Phd2ko* and wild-type CBs and AMs is provided for all annotated genes detected in a separate Excel resource.

**Supplementary Table 2. Selection of genes from Supplementary Table 1 that are, or have been implicated to be, involved in oxygen chemosensitivity.** Provided in a separate Excel resource.

**Supplementary Table 3. Gas composition in test solutions used for tissue superfusion in fluorescent imaging experiments.** For details of fluorescent imaging protocol, see the main manuscript. 50 µM doxapram solution was prepared by diluting a 1000x doxapram hydrochloride (USP) stock in DMSO in control solution. Anoxic solution was supplemented with 4 mg·ml$^{-1}$ sodium sulphite (Merck).

| Test solution | Compressed air | $N_2$ | $CO_2$ |
|---:|:---:|:---:|:---:|
| Control,<br>50 µM doxapram,<br>45 mM $K^+$ | 95% | 0% | 5% |
| 10% $O_2$ | 48% | 47% | 5% |
| 5% $O_2$ | 23% | 72% | 5% |
| 2% $O_2$ | 9% | 86% | 5% |
| 1% $O_2$<br>Anoxia | 2% | 93% | 5% |
| 10% $CO_2$ | 90% | 0% | 10% |

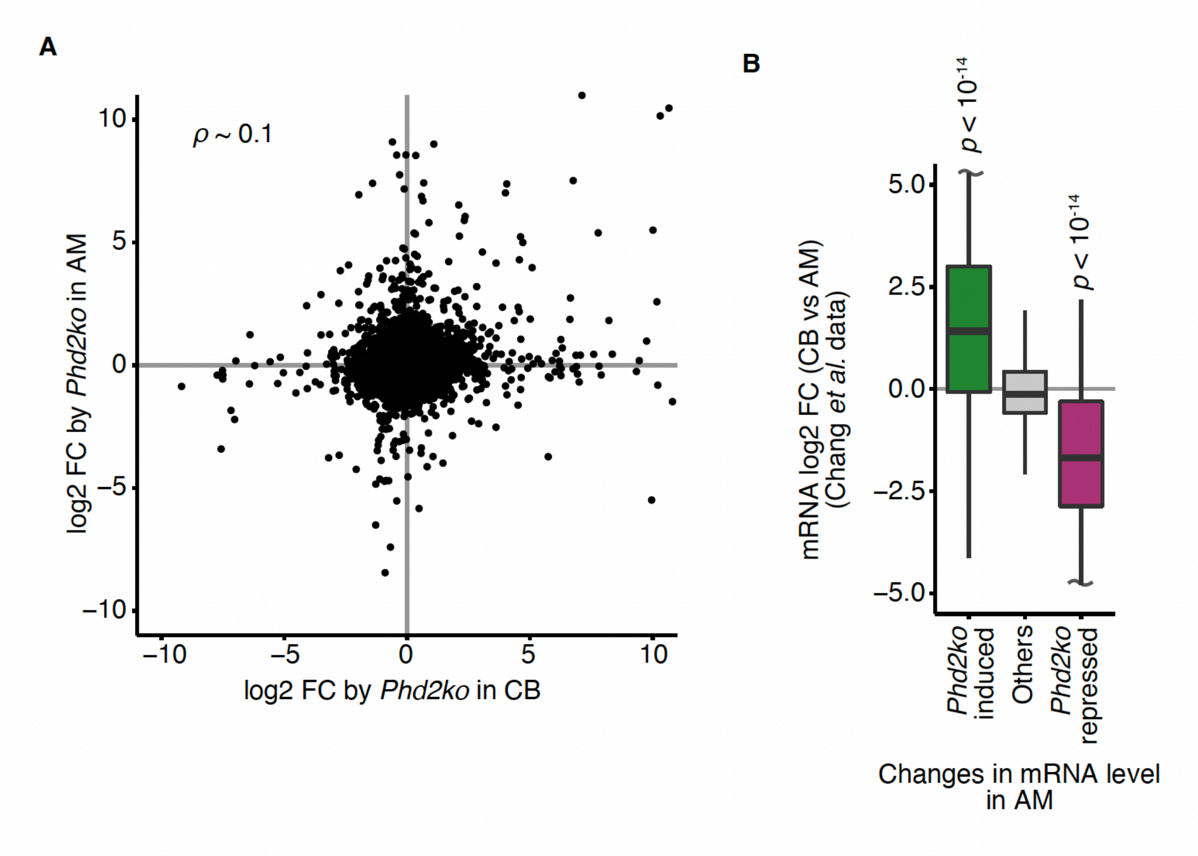


**Supplementary Figure 1. RNA Sequencing of *Phd2ko* (*Phd2*$^{f/f}$*;THCre*) versus wild-type (*Phd2*$^{f/f}$) adrenal medullas (AMs) and carotid bodies (CBs).** (**A**) Comparison of the effects of *Phd2ko* on mRNA abundance in the carotid body (CB) with that in the adrenal medulla (AM). The correlation was analysed with the Spearman's rank coefficient (n = 9,867). Genes with too little expression (transcript per million ≤ 10 in all of the conditions analysed) were excluded from the analysis. (**B**) Similar to Figure 1C (main text), but the fold difference of mRNA abundance between the CB and AM was calculated using the data from (Chang et al. 2015) instead of data generated in this paper. The distribution for genes induced (green) or repressed (purple) by *Phd2ko* in the AM was compared against that for all other genes (grey) using the two-sided Mann-Whitney U test (n = 114, 80, and 15,315, respectively).

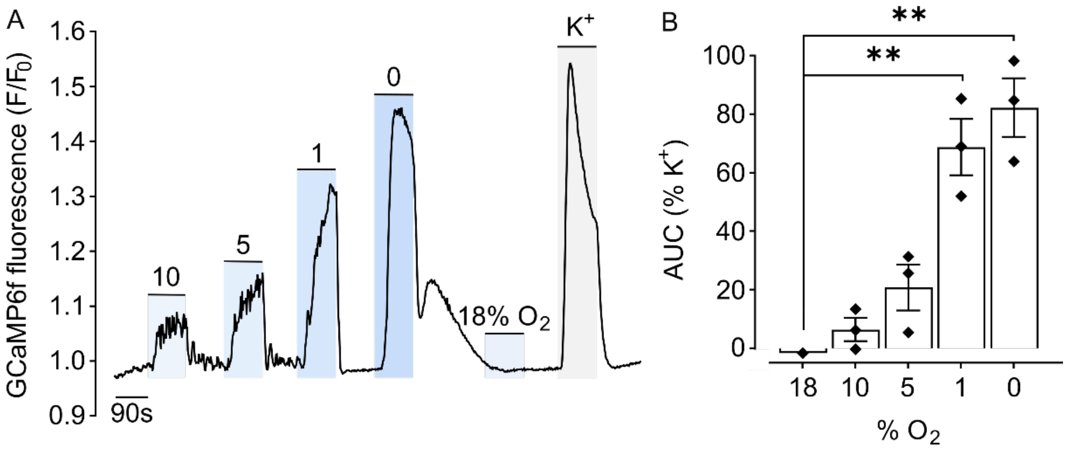


**Supplementary Figure 2. $Ca^{2+}$ imaging showing acute oxygen sensitivity in the carotid body from mice expressing GCaMP6f in $TH^+$ cells (*$Ai95^{f/+}$;THCre* or 'wild-type' mice).** (**A**) Representative trace of $F/F_0$ GCaMP6f fluorescence in the $K^+$ responsive region of the carotid body exposed to various stimuli (denoted in blue/grey shaded areas): 10, 5, 1% $O_2$, anoxia, 45 mM $K^+$ or a sham (18% $O_2$) control. (**B**) Responses to hypoxia in the carotid body are quantified as the average area under curve, presented as % of the response to high $K^+$. Mean ± S.E.M. are plotted with individual values shown. Data were analysed by a one-way ANOVA, $P$ = 0.0005, followed by Dunnett's multiple comparisons test, ** $P < 0.01$.

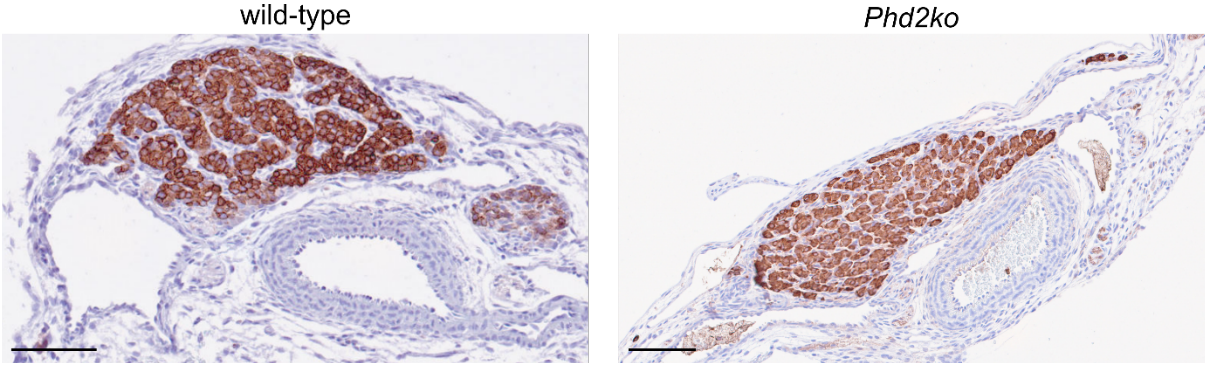


**Supplementary Figure 3. Organ of Zuckerkandl (OZ) in neonatal wild-type (*Phd2$^{f/f}$*) and *Phd2ko* (*Phd2$^{f/f}$;THCre*) mice.** Representative images showing chromogranin A (CgA) immunohistochemistry (brown) in the OZ in neonatal (P0.5) wild-type and *Phd2ko* mice. Scale bars show 0.1 mm.

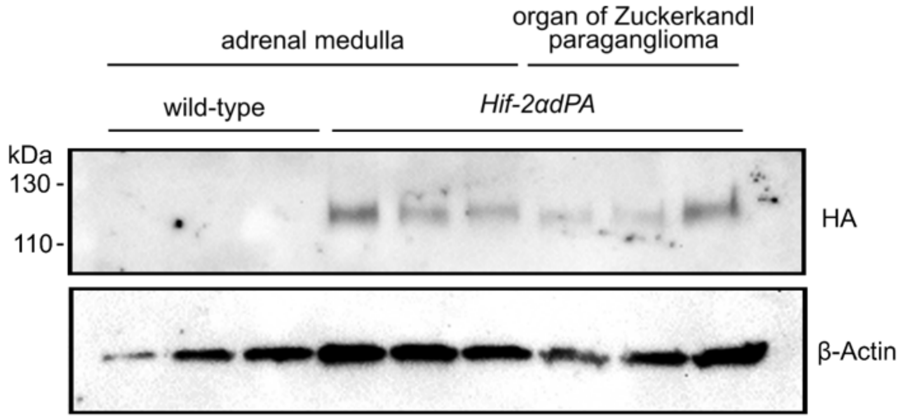


**Supplementary Figure 4. HIF-2α protein stabilisation in chromaffin tissues of *Hif-2αdPA* (*Hif-2αdPA$^{f/+}$;THCre*) versus wild-type mice.** Immunoblot for hemagglutinin (HA) or β-actin confirming HA-tagged *Hif-2αdPA* transgene expression in the adrenal medulla (AM) and organ of Zuckerkandl (OZ) from *Hif-2αdPA*, but not wild-type, mice (n = 3 mice for AM and OZ, with one mouse per lane).

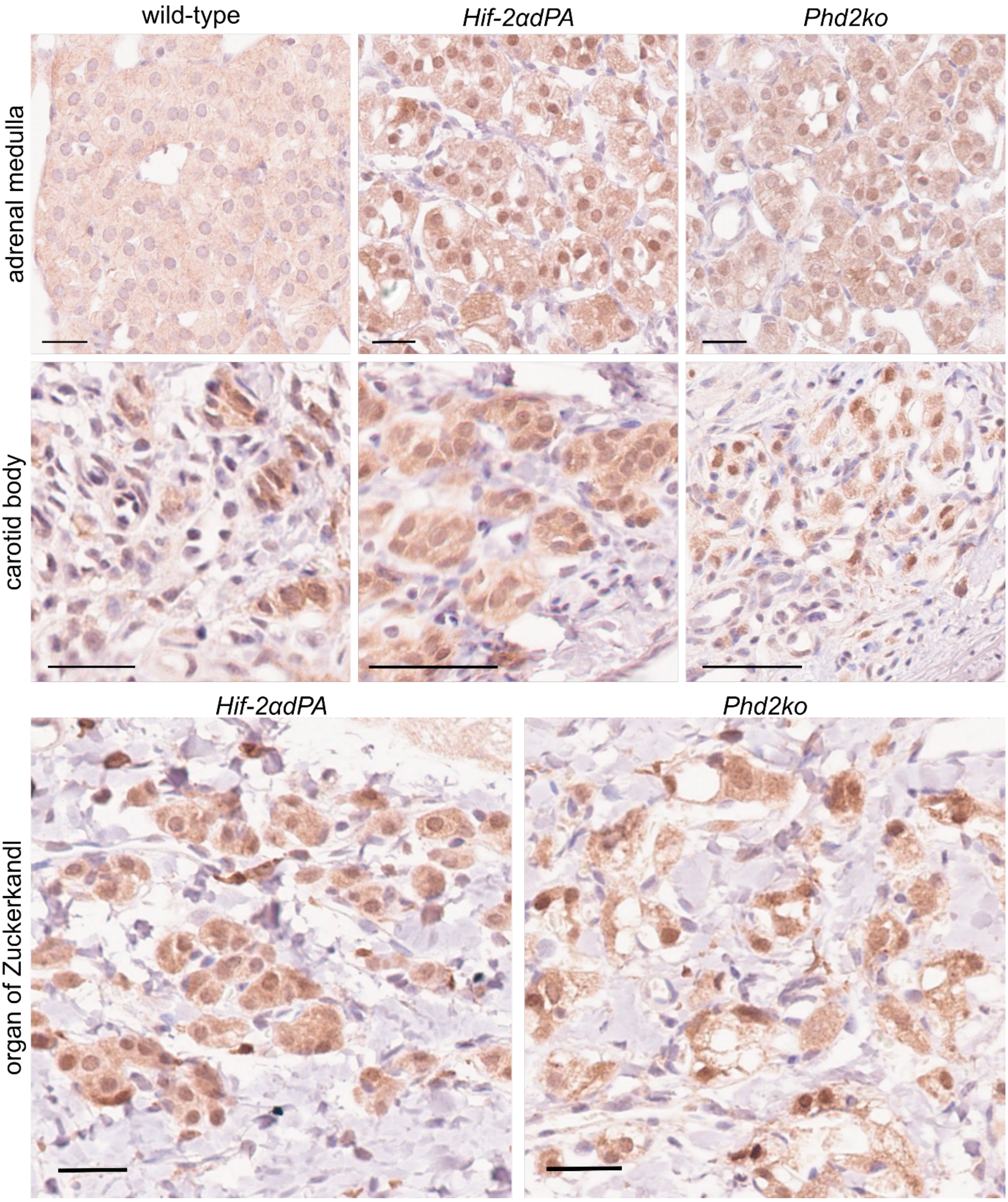


**Supplementary Figure 5. HIF-2α protein stabilisation by immunohistochemistry in chromaffin tissues of *Hif-2αdPA* (*Hif-2αdPA$^{f/+}$;THCre*) and *Phd2ko* (*Phd2$^{f/f}$;THCre*) mice.** HIF-2α immunohistochemistry showing nuclear HIF-2α protein stabilisation in chromaffin cells from the adrenal medulla (top row), carotid body (second row) and organ of Zuckerkandl (bottom row) of *Hif-2αdPA* (middle) and *Phd2ko* (right), but not wild-type (left, where present), adult mice. Scale bars represent 0.025 mm.

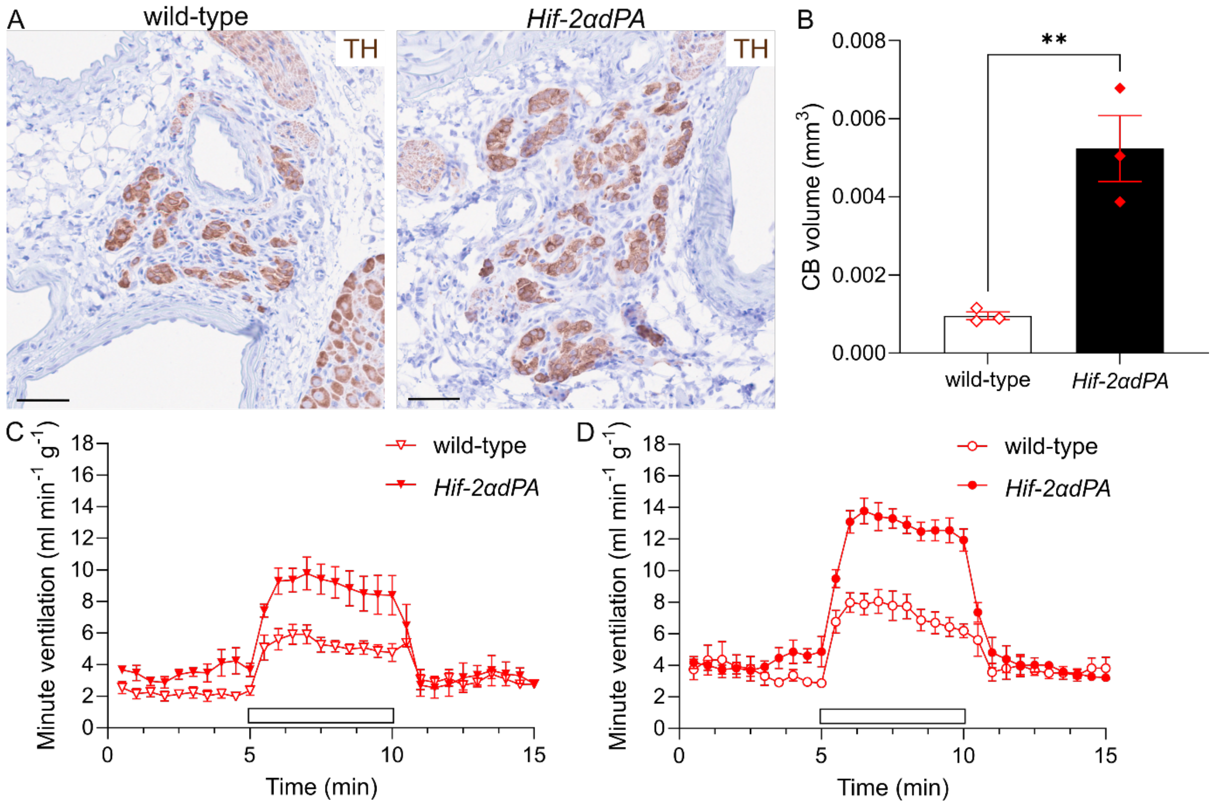


**Supplementary Figure 6. Paraganglioma-like CBs and enhanced hypoxic ventilatory responses in *Hif-2αdPA* (*Hif-2αdPA$^{f/+}$;THCre*) compared to wild-type mice.** (**A**) Tyrosine hydroxylase immunohistochemistry (TH)(brown) in CB sections from wild-type and *Hif-2αdPA* mice. Scale bars show 0.05 mm. (**B**) CB volume calculated from the area of $TH^+$ structures in sequential sections in adult wild-type and *Hif-2αdPA* mice. Bars show mean ± S.E.M. with individual values displayed. Data were analysed by an unpaired Student's *t*-test, ** $P < 0.01$. (**C, D**) Hypoxic ventilatory responses to 10% $O_2$/3% $CO_2$ (white box) in (**C**) male (n=5) and (**D**) female (n=6) *Hif-2αdPA* and wild-type mice. Acute response to hypoxia, quantified as the difference between minute ventilation in the first minute of stable hypoxia and the last minute of room air is 8.6 ± 0.5 ml·min$^{-1}$·g$^{-1}$ in *Hif-2αdPA* versus 5.0 ± 0.6 ml·min$^{-1}$·g$^{-1}$ in wild-type females (mean ± S.E.M., unpaired Student's *t*-test $P$ = 0.0008) and 5.4 ± 0.5 ml·min$^{-1}$·g$^{-1}$ in *Hif-2αdPA* versus 3.6 ± 0.4 ml·min$^{-1}$·g$^{-1}$ in wild-type males ($P$ = 0.029).

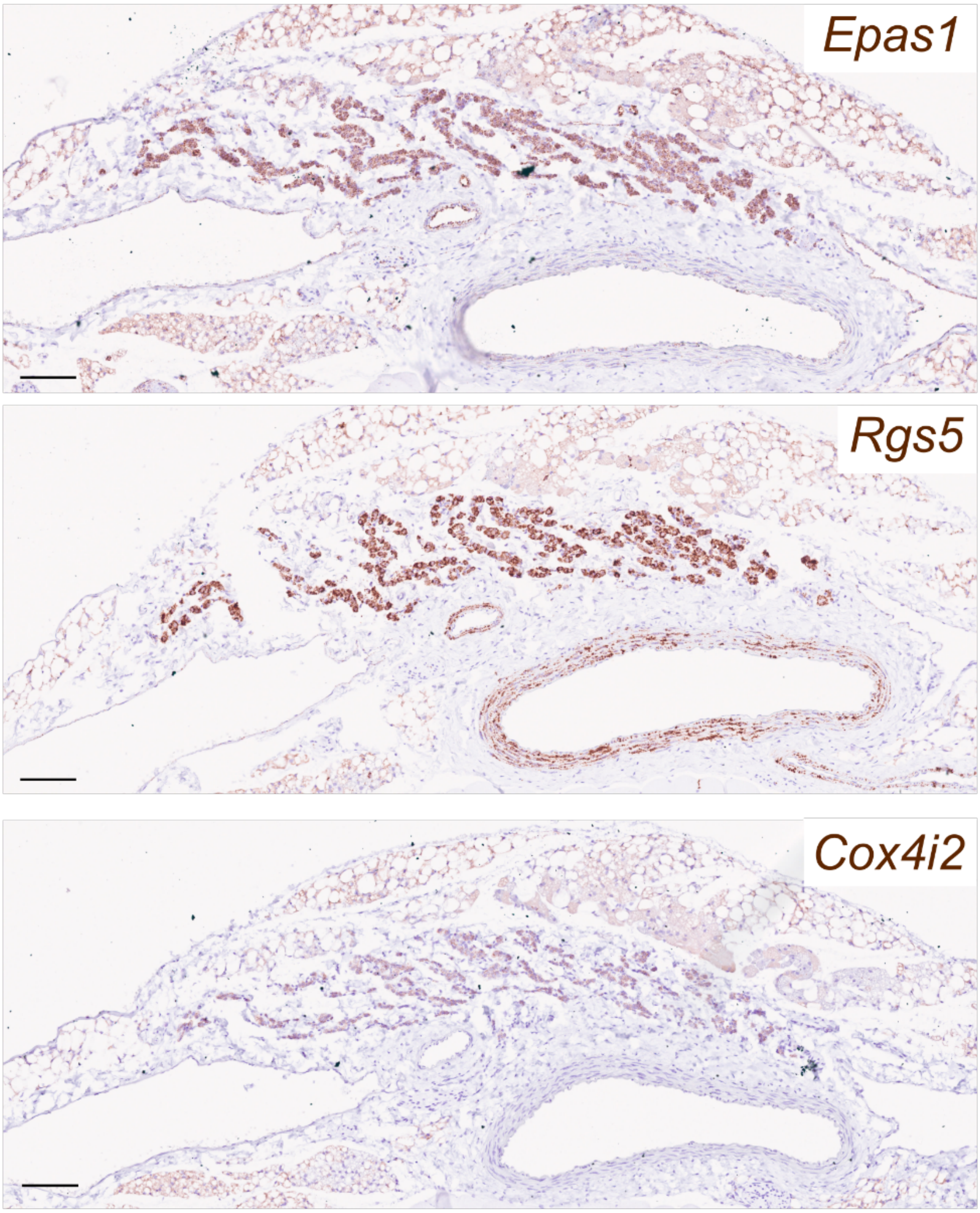


**Supplemental Figure 7. Gene expression in *Hif-2αdPA* (*Hif-2αdPA*$^{f/+}$*;THCre*) abdominal organ of Zuckerkandl paraganglioma (OZ PGL).** Representative images of *in situ* hybridisation for *Epas1, Rgs5* and *Cox4i2* mRNA (brown) in the OZ PGL from an adult *Hif-2αdPA* mouse. Scale bars are 0.1 mm.
